# Graph combinatorics based group-level network inference

**DOI:** 10.1101/758490

**Authors:** Shuo Chen, Qiong Wu, L. Elliot Hong

**Affiliations:** Division of Biostatistics and Bioinformatics and Maryland Psychiatric Research Center, University of Maryland School of Medicine; Department of Mathematics, University of Maryland, College Park; Maryland Psychiatric Research Center, Department of Psychiatry, University of Maryland School of Medicine

**Keywords:** combinatorics, graph theory, network topology, *l*_0_ norm regularization, network statistics

## Abstract

We consider group-level statistical inference for networks, where outcomes are multivariate edge variables constrained in an adjacency matrix. The graph notation is used to represent a network, where nodes are identical biological units (e.g. brain regions) shared across subjects and edge-variables indicate the strengths of interactive relationships between nodes. Edge-variables vary across subjects and may be associated with covariates of interest. The statistical inference for multivariate edge-variables is challenging because both localized inference on individual edges and the joint inference of a combinatorial of edges (network-level) are desired. Different from conventional multivariate variables (e.g. omics data), the inference of a combinatorial of edges is closely linked with network topology and graph combinatorics. We propose a novel objective function with 𝓁_0_ norm regularization to robustly capture subgraphs/subnetworks from the whole brain connectome and thus reveal the latent network topology of phenotype-related edges. Our statistical inferential procedure and theories are constructed based on graph combinatorics. We apply the proposed approach to a brain connectome study to identify latent brain functional subnetworks that are associated with schizophrenia and verify the findings using an independent replicate data set. The results demonstrate that the proposed method achieves superior performance with remarkably increased replicability.

## 1 Introduction

Modeling group-level network data has been an active area of research in statistics. For example, brain connectome research often aims to investigate whether brain functional and/or structural networks are associated with behavioral and symptomatic phenotypes, and microbiome network studies focus on whether microbial networks are influenced by the clinical status (Lukemire et al., 2017; Xia and Li, 2017; Cai et al., 2018; Simpson et al., 2019; Warnick et al., 2018; Shaddox et al., 2018). In these applications, the data structure of each subject can be represented by a graph, where a node represents a well-defined biological unit (e.g. a brain area) and an edge indicates the interactive relationship between a pair of nodes. We consider all nodes are identical across subjects because brain regions are shared across all human subjects.

The outcome variables ofgroup-level network data are edge-variables quantifying the strengths of interactions between nodes, which vary across subjects and can be associated with external phenotypes (e.g. the clinical diagnosis and treatment response). In that, edges are (weighted/continuous or binary) multivariate outcomes that are constrained by a set of nodes in an adjacency matrix. The statistical inferential procedure of multivariate edge-variables is different from conventional high-throughput statistical methods, for example, the commonly used false positive discovery rate (FDR) and shrinkage method (Benjamini and Hochberg, 1995; Efron et al., 2004) because edges are intrinsically linked with network topology and graph combinatorics. Directly applying multivariate edgewise inference often leads to a high false positive discovery rate and more importantly, low interpretability, because individual edges can not reveal the systematic influence of the phenotype at a network level (Xia et al., 2019). On the other hand, comparing graph summary statistics like ‘small-worldenss’ and betweenness loses the spatial specificity (Craddock et al., 2013). Therefore, our goal of group-level network inference is to extract each unknown subnetwork that is a combinatorial set of phenotype-related edges covered by an organized topological structure, and provide both network and edge level inference. To address this challenge, group-level network analysis methods have been developed using advanced statistical techniques (Kim, 2014; Narayan et al., 2015; Ginestet et al., 2017; Zhang et al., 2017; Durante et al., 2018; Xia and Li, 2018; Kundu et al., 2018; Wang et al., 2019; Higgins et al., 2018; Cao et al., 2019 among others). These methods have greatly improved the accuracy of statistical inference and yield numerous meaningful biological findings.

In this current research, we propose a new statistical framework for group l**e**vel network inference (GLEN) consisting of two steps (Figure 1). Firstly, latent subnetwork extraction: GLEN implements a novel *l*_0_ norm shrinkage based optimization algorithm to extract subgraphs that cover the maximal number of phenotype-related edges by the minimum size (i.e. minimizing the number of edges in the subgraphs). We define a subnetwork as an induced subgraph with an organized topological structure (e.g. community) in the graph/network space (here ‘subnetwork’ and ‘subgraph’ are exchangeable). Next, we perform statistical inference based on graph combinatorics and network topology to produce results with higher power and lower false positive discovery rates, and meaningful biological interpretability. Both network and edge level inference can be obtained. The property of graph combinatorics is the foundation for the two-step GLEN method because the *l*_0_ norm shrinkage algorithm can only detect subnetworks with extreme low combinatorial probability in a random graph model and the graph combinatorial probability is directly used for statistical inference. In addition, GLEN can also be a complement to the existing methods. For example, the locations and topological structures of phenotype-related subgraphs detected by GLEN can become prior information for existing network analysis models (Xia and Li, 2017; Simpson et al., 2019; Xia et al., 2019).

**Figure 1:**
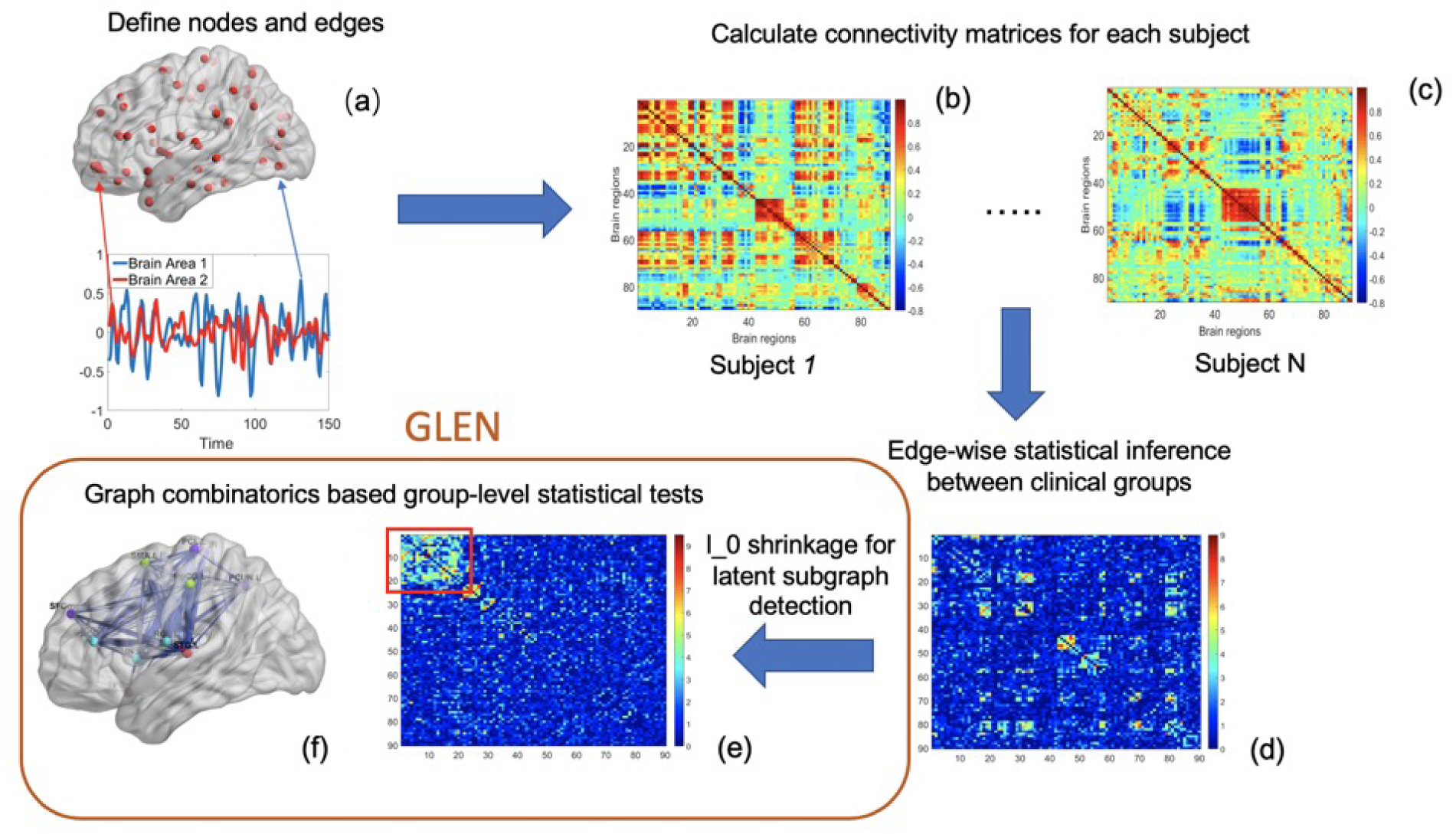
a) Define brain regions as nodes and connectivity metrics between each pair of nodes as edges. b c) Calculate the connectivity matrix for each single subject in a study where each off-diagonal element in the matrix represents the connectivity strength between two nodes; then identify differential connectivity patterns between clinical groups. d) Plot the edge-wise statistical inference where each off-diagonal element is a log transformed pvalue (e.g. two sample test p-value per edge between clinical groups and a hotter point in the heatmap suggests larger group-wise difference). e) Reveal the disease related subnetwork detected by GLEN. f) Shows the corresponding 3D brain image. Note that e) is obtained by re-ordering the nodes in d) by listing detected subnetwork first (i.e. these two graphs are isomorphic).

We apply the proposed method to an example data of brain connectome study for schizophrenia research with a primary data set and an independent validation data set. The proposed method along with comparable methods are applied to both data sets separately. We find that the subnetwork identified by GLEN in the primary data set is almost identical to the validation data set, and thus the findings are highly replicable. In contrast, the comparable methods only detect none or a small proportion of edges that are shared by both data sets. These findings are further supported by our simulation results that the false positive and false negative discovery rates are simultaneously reduced by GLEN when phenotype related edges consist of dense subgraphs with organized topological structures. The biological findings using GLEN may also provide novel insights into neurophysiology and neuropathology. For example, GLEN can detect that the interconnections between three well-known brain networks (the default mode network, executive network, and salience network) are associated with a brain disease.

## 2 Methods

### 2.1 Background

We consider a group of networks 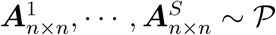, where *S* is the number of subjects and all networks share an identical graph representation *G* = {*V, E*} with |*V*| = *n* nodes and |*E*| = *n*(*n* − 1)*/*2 edges. For subject *s*, the multivariate outcome 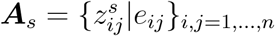 (*e*_*ij*_ denotes an edge connecting nodes *i* and *j*) can be a binary or weighted adjacency matrix and a vector of covariates ***x***_*s*_ are also observed (e.g. clinical and demographic variables).

We assume that ***A***_*s*_ follows a distribution 𝒫 with parameters related to ***x***_*s*_.

We let 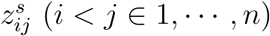 denote an off-diagonal edge entry in ***A***_*s*_, and let 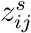 follow a conditional distribution of the exponential family (e.g. a Bernoulli distribution for the binary adjacency matrix ***A***_*s*_ and a Gaussian distribution for the weighted adjacency matrix ***A***_*s*_, Bowman, 2005; Derado et al., 2010; Risk et al., 2018):

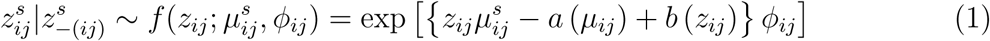

where 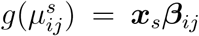. Recently, advanced statistical methods have been developed to provide localized (edge-wise) statistical inference while accounting for the dependence between edges for group-level network inference (Xia and Li, 2017; Xia and Li, 2018; Chen et al., 2018). These methods often yield improved statistical inferential results (e.g. testing statistics and p-values) on individual edges by taking into account the covariance structure between edges. However, the results of these methods still face the issue of multiple testing for multivariate edges to identify *E*_1_ = {*e*_*ij*_|***β***_*ij*_ ≠ 0} ⊂ *E*. If edges in *E*_1_ are randomly distributed in whole brain connectome *G*, conventional methods for multivariate statistical inference (e.g. FDR and FWER) are applicable to the *n*(*n* − 1)*/*2 vector because the order of edges can be randomly shuffled with no impact on the results. However, in practice, phenotypes rarely influence brain connections (edges) that are randomly distributed in the brain, instead, most times systematically.

In this paper, we focus on group-level network statistical inference which statistically assesses the impact of ***x***_*s*_ on ***A***_*s*_ at the subnetwork level. We consider *G*_*c*_ ⊂ *G* as a subnetwork with high density of phenotype-related edges, where Σ*I*(*β*_*ij*_ ≠ 0|*e*_*ij*_ ∈ *G*_*c*_)*/*|*E*_*c*_| >> |*E*_1_|*/*|*E*|. Since edges in *E*_1_ are not randomly distributed in *G, G*_*c*_ may have an organized topological structure. We define the lower bound of the subnetwork size (see subsection 2.3). The underlying topological structures of {*G*_*c*_} are inherently related to the graph theory and combinatorics which has a significant impact on network level inference. However, in practice, both *E*_1_ and the subnetworks {*G*_*c*_} are latent and often overwhelmed by false positive noise.

#### Input Data of GLEN

We denote the resulting edge-wise statistical inference matrix as **W**_0_, where 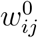 is an entry in **W**_0_ (e.g. test statistics *t*_*ij*_ and p values − log(*p*_*ij*_)). We first perform Sure Independence Screening (SIS) on **W**_0_ (Fan and Lv, 2008). We denote screened matrix **W**, where the off-diagonal entry 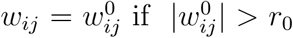 and *w*_*ij*_ = 0 if 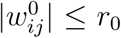, where *r*_0_ is the cutoff. **W** can effectively exclude most non-informative false positive edges while maintaining a high proportion of true positive edges (Fan and Lv, 2008; Li et al., 2012). We consider **W** and the graph notation *G* = {*V, E*, **W**} as the input data of our method. The goal of our analysis is to accurately identify the true phenotype related latent subnetworks that are maximally composed of edge set *E*_1_.

### 2.2 Detecting subgraphs via 𝓁_0_ norm regularization

If *G* = {*V, E*, **W**} is a non-random graph, then there exist subgraphs 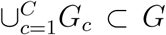 and *G*_*c*_ = {*V*_*c*_, *E*_*c*_} with higher density of phenotype-related edges than the rest of *G*. In other words, edges in *G*_*c*_ are more likely to be non-null: ℙ(*β*_*ij*_ ≠ 0|*e*_*ij*_ ∈ *G*_*c*_) > ℙ(*β*_*ij*_ ≠ 0|*e*_*ij*_ ∉ *G*_*c*_), which can be reflected by 𝔼(*w*_*ij*_|*e*_*ij*_ ∈ *G*_*c*_) > 𝔼(*w*_*ij*_|*e*_*ij*_ ∉ *G*_*c*_) from the input data **W**. We provide details of statistically testing whether *G* = {*V, E*, **W**} is a (weighted) random graph in the supplementary materials.

Our goal is to extract subnetworks {*G*_*c*_} that 𝔼(*w*_*ij*_|*e*_*ij*_ ∈ *G*_*c*_) ≫ 𝔼(*w*_*ij*_|*e*_*ij*_ ∉ *G*_*c*_) and 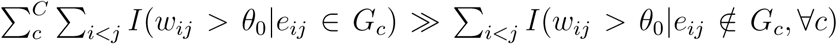 where *θ*_0_ is a reasonable threshold. The estimand subgraphs {*G*_*c*_} can be linked with input data **W** by a subgraph based matrix 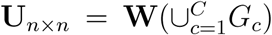. The block diagonal matrix 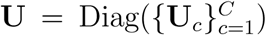 or 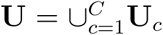 is determined by the subgraphs 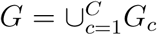, where an entry *u*_*ij*_ = *w*_*ij*_ if the corresponding edge *e*_*ij*_ ∈ *G*_*c*_, ∀*c* and *u*_*ij*_ = 0 if *e*_*ij*_ ∉ *G*_*c*_. *G*_*c*_ can be a singleton that *V*_*c*_=1 and |*E*_*c*_| = 0. Thus, a diagonal submatrix **U**_*c*_ bijectively corresponds to a subgraph *G*_*c*_ and **W**. Since *G*_*c*_ often shows an organized topological structure, we further denote the latent organized topological pattern of *G*_*c*_ by 𝒯(*G*_*c*_). Our goal is to identify phenotype related subnetworks/subgraphs 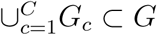 from **W**, and to learn 𝒯 (*G*_*c*_).

**W** is often less dense (e.g. 5 ∼ 20% edges are non-zero entries after screening). The existing algorithms for subgraph/subnetwork detection (e.g. community detection) can be substantially impacted due to the less dense network structure and false positive edges (Newman and Girvan, 2004 and Rohe et al., 2011). This could be more challenging for a weighted matrix **W**. To achieve our goal above, we propose a novel 𝓁_0_ norm regularization based objective function that minimizes the sizes of the phenotype-related topological structures (subgraphs) 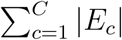 and thus suppresses the impact of random false positive noise and better reveal the latent subnetworks.

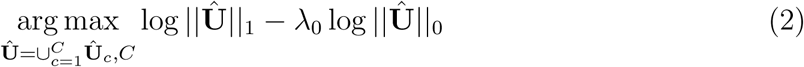

where 1 ≤ *C* ≤ *n*, and in 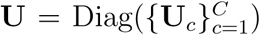 each **Û**_*c*_ corresponds to a subgraph *G*_*c*_ = {*V*_*c*_, *E*_*c*_} that 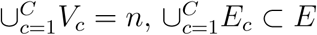, and *V*_*c*_ ∩ *V*_*c′*_ = ∅.

We maximize the first term ‖**Û**‖_1_ = Σ_*c*_ Σ_*i*<*j*_ (*w*_*ij*_|*e*_*ij*_ ∈ *G*_*c*_) to cover more high-weight edges in **W** by the set of subgraphs 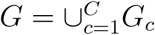 (*G*_*c*_ is clique by default). Maximizing the first term can reduce the chance to miss the true positive edges. The second term is defined by 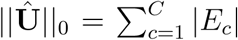, which is a penalty term. Generally, the first term increases when the size ‖**Û**‖_0_ is larger because a greater ‖**Û**‖_0_ can cover more edges. However, a larger size of ‖**Û**‖_0_ tends to include more false positive edges. The objective function aims to maximize true positive findings while penalizing the 𝓁_0_ norm of **Û** (false positive findings).

#### Tuning parameter

*λ*_0_ is a tuning parameter and can be selected using a likelihood based method (see Supplementary Materials). When default *λ*_0_ = 0.5, the proposed objective function is equivalent to the well-known problem of dense subgraph discovery in the field of graph theory and computer science and thus can be conveniently computed by linear programming and greedy algorithms (Tsourakakis et al., 2013; Miyauchi and Kakimura, 2018). The density of *G*_*c*_ is defined by 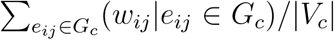 and the density in *G*_*c*_ is higher than all other possible induced subgraphs (Gionis and Tsourakakis, 2015). However, the group level network inference for our applications may involve more complex scenarios because a set of subgraphs with unknown sizes and topological structures, and thus the existing dense subgraph discovery algorithms may not be directly applicable. Therefore, we resort it to the 𝓁_0_ norm regularization for subgraph extraction.

#### 𝓁_0_ *norm regularization*

The 𝓁_0_ norm regularization has been attracting interest in the field of statistics (Shen et al., 2012; Li et al., 2018; Hazimeh and Mazumder, 2018). Our optimization is distinct from these methods because we focus on edge-variables in an adjacency matrix. Interestingly, we note that the penalty term ‖**Û**‖_0_ is related to the number of subgraphs *C*, which links the network/graph detection with 𝓁_0_ norm regularization. This property is unique for edge-variables in a graph space and becomes the foundation for computationally efficient heuristic of 𝓁_0_ norm regularization for edge variables. The increase of *C* generally leads to smaller sizes of organized topological structures, and thus reduced 𝓁_0_ norm but larger loss of ‖**Û**‖_1_. If *C* = *n* then ‖**Û**‖_0_ = 0, and *C* = 1 then ‖**Û**‖_0_ = *n* × (*n* − 1)*/*2.

To illustrate this point, the objective function can also be re-written as (similar to the counterpart of expression for LASSO):

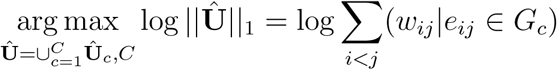

with subject to

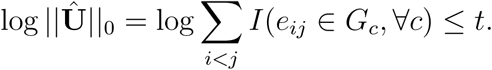

We note that *C* can provide an upper bound for ‖**Û**‖_0_.

##### Lemma 1.

*For a given value of C, we have the upper bound* sup ‖***Û***‖_0_ = (*n* − *C* + 1)(*n* − *C*))*/*2.

*Proof*. For *C* = 2,

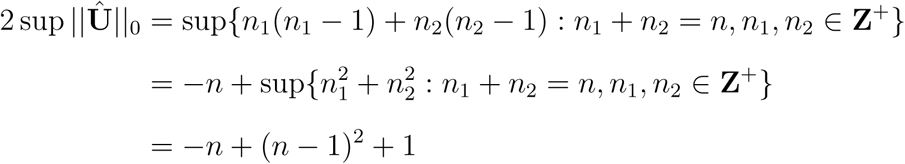

where *n*_1_ = |*V*_1_|, *n*_2_ = |*V*_2_| are number of vertices for the two communities. Hence, 2 sup ‖**Û**‖_0_ = (*n* − *C* + 1)(*n* − *C*) for C=2.

Inductively, assume it’s true for *C* = *k* −1, such that 2 sup ‖**Û**‖_0_ = (*n*−*k* +2)(*n*−*k* +1). Then, for *C* = *k*,

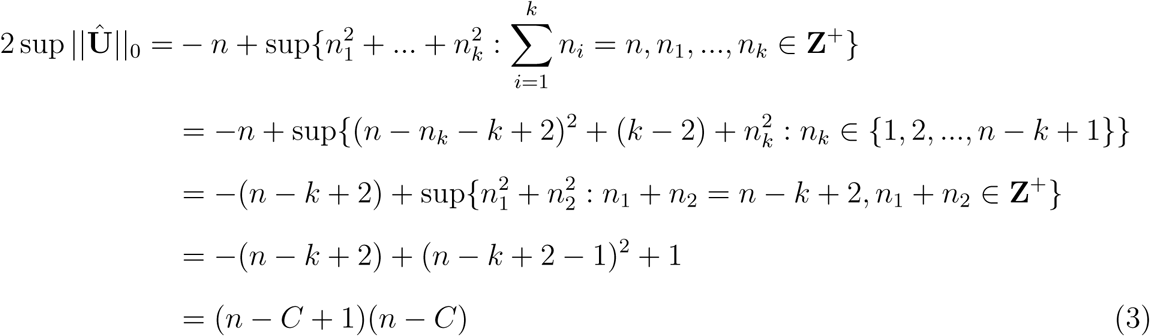

Therefore, the result is true for *C* = *k*. Hence, the claim is proved. □

Lemma 1 shows that increasing *C* can effectively shrink the size of ‖**Û**‖_0_. We implement the path to shrink ‖**Û**‖_0_ by increasing *C*. Specifically, the algorithm first optimizes the objective function (2) with a given *C* and the corresponding upper bound of ‖**Û**‖_0_ = (*n* − *C* + 1)(*n* − *C*) to identify subgraphs covering maximal edge-weights. Then, we exhaust *C* from 2 to *n* − 2 and perform grid search to estimate *Ĉ*. The combination of *Ĉ* and **Û** that optimize (2) become the final estimates.

The 𝓁_0_ norm regularization is critical to suppress false positive noise. Let *w*_*ij*_ ≠ 0|*β*_*ij*_ = 0 be an false positive edge that *i* ∈ *G*_*c*_ and *j* ∉ *G*_*c*_, ∀*c*. Adding *e*_*ij*_ to *G*_*c*_ will increase ‖**Û**‖_0_ by |*V*_*c*_|. However, the increase of ‖**Û**‖_1_ is small because there is only a small proportion of *w*_*i′j*_ > 0, *i*′ ∈ *G*_*c*_ and *i*′ ≠ *i*. In another scenario, *w*_*ij*_ ≠ 0|*β*_*ij*_ = 0 be an false positive edge that *i* ∈ *G*_*c*_ and *j* ∈ *G*_*c*′_, ∀*c* ≠ *c*′. The inclusion of a false positive edge *e*_*ij*_ can connect two subgraphs false positively and increase ‖**Û**‖_0_ by |*V*_*c*_| × |*V*_*c′*_ |. In both cases, the inclusion of false positive edges leads to a high cost for the pentalty term. Therefore, the 𝓁_0_ norm regularization is particularly useful to extract informative subgraphs from a noisy and less dense matrix **W**.

The details of the algorithm, the derivation, and the software package for the objective function are provided in the Supplementary Materials.

#### Consistency for subgraph detection

In the following Lemma 2 and Theorem 1, we establish that for given true number of subgraphs *C*\*, the proposed objective function can provide a consistent estimate for the community topological structure (the collection of node-induced subgraphs) in the sense that the error of assignments for nodes is negligiable in large graphs. Furthermore, considering that *C* is optimized by grid search, the estimated community structure based on the optimal number of subgraphs selected by objective function (2) also yields a consistent estimator.

To formalize the theoretical results, in a graph representation of input **W**, we assume there exists a set of subgraphs 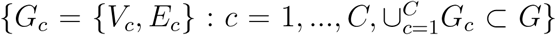 for phenotype-related edges, such that 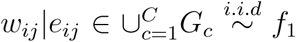 and 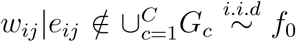 where *f*_1_ and *f*_0_ are continuous densities in [0, 1] with mean and variance 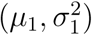 and 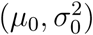, *μ*_1_ > *μ*_0_, respectively. Then the expectation matrix ***P*** = 𝔼(**W**) can be expressed as ***P*** = **Θ*B*Θ**^*T*^, where **Θ** ∈ ℝ^*n*×*C*^ is a membership matrix indicating the community index of each node, and ***B*** ∈ ℝ^*C*×*C*^ equals *µ*_1_ in diagonal and *µ*_0_ for others.

For convenience, we use notation

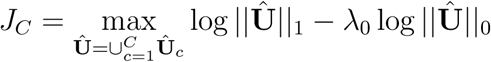

which is the maximized objective function with *C* subgraphs. Then the objectivie function (2) yields a solution 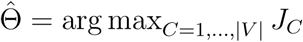. Let Θ* be the true matrix of membership.

##### Lemma 2

(Consistency with known *C*\*). *Assume* ***P*** *of rank C has smallest absolute nonzero eigenvalue ξ and* 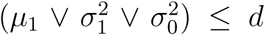 *for d* ≥ *c*_0_ log *n and c*_0_ > 0. *Then, if* 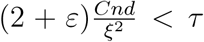 *for some τ, ε* > 0, *the output* 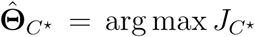 *is consistent up to a permutation. Equivalently, if Ŝ*_*c*_ *is the estimated nodes set for subgraph G*_*c*_, *c* = 1, …, *C*\*. *Then Ŝ*_*c*_ ∩ *V*_*c*_ *is the set in V*_*c*_ *that the assignment of nodes can be guaranteed, and with probability at least* 1 − *n*^−1^, *up to a permutation, we have*

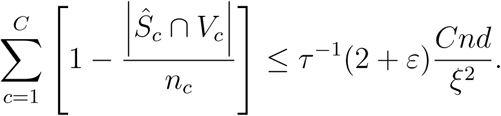

*with n*_*c*_ = |*V*_*c*_|, *c* = 1, …, *C*\*.

##### Theorem 1

(Consistency for grid searched *C*). *Let the sizes of subgraphs n*_*c*_, *c* = 1, …, *C*\* *be generated from a multinomial distribution with probabilities* ***π*** = (*π*_1_, …, *π*_*C*_*\**). *Assume* ∃*δ* > 0, *such that*

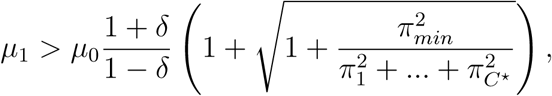

*then under conditions in Lemma 2 and λ*_0_ = 1*/*2, *the number of mis-assigned nodes satisfy n*_*e*_ = *o*_*p*_(*n*_min_) *as n* → ∞ *where n*_min_ *is the size of the smallest subgraph*.

Proof. The proofs of Lemma 2 and Theorem 1 are included in the supplemental materials. Note that the theorem is also true for a general setting of weighted stochastic block model with 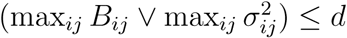.

*Extracting complex topological structures* 𝒯(*G*_*c*_) *by the objective function (2)*: we consider the community/clique structure as the default topological structure, although the objective function (2) can be further optimized if the phenotype related edges can be covered more subgraphs with more sophisticated topological structures. For example, there are two potential subgraphs, one is a clique *G*_*c*_ and the other is a bipartite subgraph *G*_*c′*_ with the same number of nodes *V*_*c*_ = *V*_*c′*_. The number of edges of the clique is greater than the bipartite subgraph *E*_*c*_ > *E*_*c′*_. If the edges with *β*_*ij*_ ≠ 0 are equivalently covered by *G*_*c*_ and *G*_*c′*_, the bipartite subgraph is preferred by (2). The detailed algorithms for more sophisticated topological structures (e.g. k-partite structure, rich-club, and interconnected subgraphs) and model selection procedure are described by Chen et al. (2019). Therefore, the complex topological structures are favored by the 𝓁_0_ norm regularization if the objective function is further optimized.

### 2.3 Graph combinatorics for network-level test

The optimization of objective function (2) reveal the underlying network topology of phenotype-related edges, which has naturally provoked graph combinatorics. The group-level network statistical inference examines whether a combinatorial set of edges (rather than individual edges) in an organized subnetwork is significantly related to the phenotype. In contrast to conventional multivariate inference on individual edges (e.g. FDR), the subnetwork based statistical inference is inherently linked with graph combinatorics as

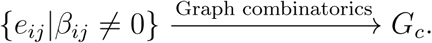

We first specify the null and alternative hypotheses of the subnetwork test. Let *ρ* = Σ_*i<j*_ *I*(*β*_*ij*_ ≠ 0)*/*|*E*| and *G* be a graph with *n* nodes and the connection probability of *ρ*. The null can be categorized into two cases: the stronger case 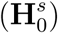 that no edge is related to the phenotype (*ρ* = 0); and the weaker case that a small proportion (*ρ* ≥ 0) of phenotype-related edges are randomly distributed in *G*. Clearly, the weaker case is more general and commonly used and the stronger case is a special case of the weaker case (Benjamini and Hochberg, 1995), and additional tests could be conducted to distinguish the two nulls as follows. Thus, we focus on the weaker case of the null. The null and alternative hypotheses are:

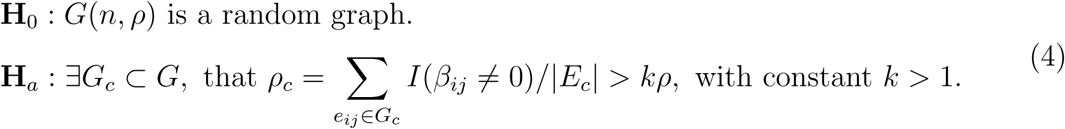

If we fail to reject the null (weaker case) that *G* is a random graph, then the conventional multi-testing methods (e.g. FDR and FWER) can be applied to identify individual edges that are related to the phenotypes. The stronger case of null can be rejected if at least one edge passed the cut-off of the multiple testing methods.

In the follows, we show that from the perspective of graph combinatorics, testing a combinatorial set of multivariate edges constrained in a subnetwork (subnetwork level inference) can lead to low false positive and false negative error rates.

We consider the permutation test for the network level inference because the asymptotic distribution of graph combinatorics based test statistic under the null is difficult (Zalesky et al., 2010; Chen et al., 2015; Chen et al., 2019). Briefly, the permutation tests generate multiple simulations (e.g. 10^4^ times) of data under the null by shuffling the labels, and calculate test statistic 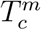 for each extracted subnetwork *G*_*c*_ and record the maximum statistic 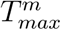 for each simulation. The percentile of observed test statistic among the maximum test statistics of all simulations becomes the *p*-value. The issues of multiple testing and selection bias can be both addressed by this procedure (Zalesky et al., 2010). We include the detailed permutation test procedure in the supplementary materials. Specifically, we assume that each simulation of the permutation test generates a matrix **W**^*m*^ under the null and the false positive edges are randomly distributed in *G* (*G* is a random graph), where *m* = 1, ⋯, *M*. The test statistic in the permutation test statistic is a function of 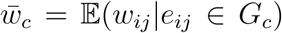 and we assume 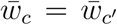. Then, the test statistic is related to the size of the subnetwork and 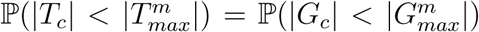 since 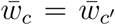. We demonstrate the impact of graph combinatorics on network-level inference in terms of Type I and Type II errors.

#### 2.3.1 Type I error of network-level inference

Type I error rate is the probability to false positively identify a significant subgraph when the null is true. For subnetwork level inference, the subgraph can be false positive only if it is detected. We first consider the probability of a (latent) dense induced subgraph existing in *G* under the null. We assume a random graph 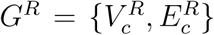 is generated with probability *p*. 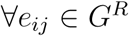, we have *δ*_*ij*_ ∼ Bern(*p*). We further let *w*_*ij*_|*δ*_*ij*_ = 1 ∼ *f*_1_ and *w*_*ij*_|*δ*_*ij*_ = 0 ∼ *f*_0_ as entries of **W**^*m*^. Therefore, the weighted adjacency matrix is generated from a random graph *G*(*n, p*) with 𝔼(*w*_*ij*_|*δ*_*ij*_ = 1) = *µ*_1_ > *µ*_0_ = 𝔼(*w*_*ij*_|*δ*_*ij*_ = 0). We prove that under the null the probability of the existence of a detectable subgraph with a density > *µ*_0_ from a random graph (null) converges to zero.

##### Theorem 2.

*For a random graph G*^*R*^ *generated as above. Let* 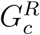 *be a detectable community structure under the null, i*.*e*.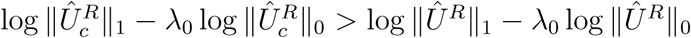, *where* 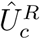 *and Û*^*R*^ *are the correspondingly subgraph-based matrix for* 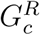 *and G*^*R*^, *respectively. Then, the existence of* 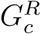 *has probability 0 asymptotically, if the number of subgraphs in* 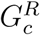 *is at most* 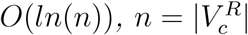, *i*.*e*.

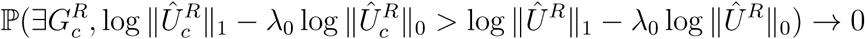

*as n* → ∞.

*Proof*. For any random graph *G*_*S*_ = (*S, E*_*S*_), let 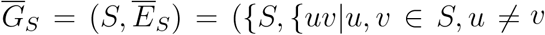, and *uv* ∉ *E*_*S*_}) be the complement of graph *G*_*S*_. Assume 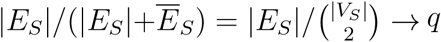, and **W**_*S*_ is the associated weighted adjacency matrix generated from the random graph with 𝔼(*w*_*ij*_|*δ*_*ij*_ = 1) = *µ*_1_ > *µ*_0_ = 𝔼(*w*_*ij*_|*δ*_*ij*_ = 0). Then, from law of large numbers, the average weights for **W**_*S*_ will have in probability

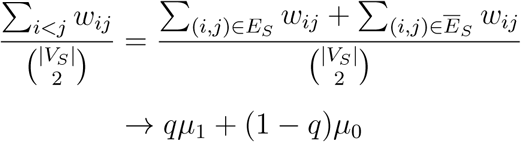

as |*S*| → ∞.

Hence, if the average weight is no less than *γ*, i.e. 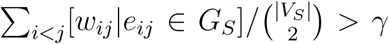, we will have asymptotically

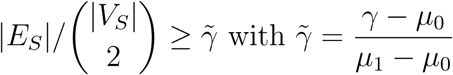

Otherwise, for any 0 < ∊ < 1, 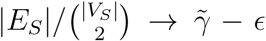, from law of large numbers, 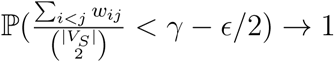 which contradicts the assumption.

Therefore, the community structure 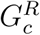 is generated from 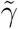-dense communities in random graph in the sense that 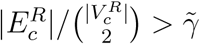. From Lemma 3 in the supplementary material A5, if 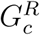 is constructed by ln *n* communities, 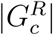 will be bounded by 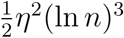 almost surely. In other words,

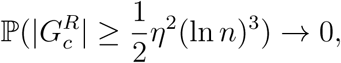

where

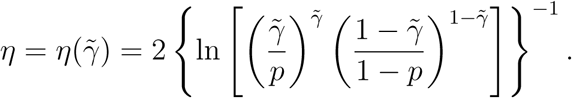

On the other hand, 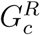 needs to be detectable community structure with respect to our objective function (2). It has been true

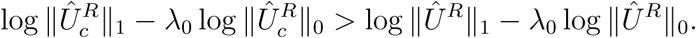

However, with probability 1 − exp(−*c/n*) for some *c* = *c*(*δ*) > 0 and *δ* > 0,

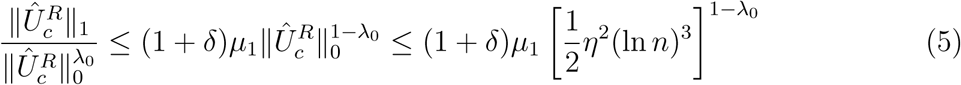

Also, with probability 1 − exp(−*c*′*/n*) for some *c*′ = *c*′(*δ*) > 0 and *δ* > 0,

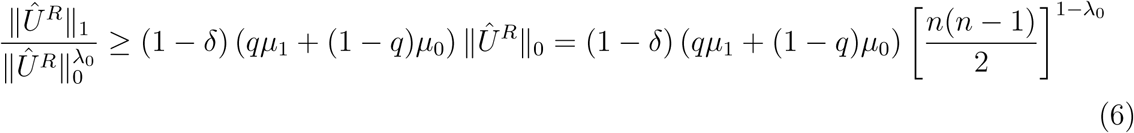

The right hand side in (5) is smaller than (6) as *n* → ∞ for *γ* ∈ (*µ*_0_, *µ*_1_) and *λ*_0_ ∈ (0, 1). Hence, our claim is true. □

A potential phenotype-related subnetwork couldn’t have larger value of objective function than the whole random graph under suitable conditions. Therefore, the probability of false positive edges comprising a detectable dense subgraph is approximately zero with a threshold level *r*_0_ under the null. The non-detectable subgraph leads to no false positive report of phenotype-related subnetwork and no Type I error for network-level inference when *H*_0_ is true.

#### 2.3.2 Type II error of network level inference

We next consider the power for network-level inference, which is the probability to reject null given that the alternative hypothesis is true and *G*_*c*_ ∈ *G* is significantly associated with the phenotype. The power is determined by the values of test statistics from all simulations of the null of the permutation test. In each simulation of the permutation test, we assume a random graph *G*^*R*^ is generated as above. Suppose *G*_*c*_ is a detectable subgraph from the data when the alternative hypothesis is true. The probability to reject the null converges to 1 when the each simulation in the permutation test yields a maximum subgraph 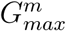 that 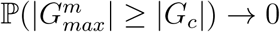 and thus 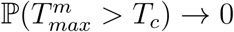.

##### Theorem 3.

*Assume that for mth simulation in the permutation test, the weighted adjacency can be regarded as a generation from random graph G*^*R*^(*n, p*) *with* 𝔼(*w*_*ij*_|*δ*_*ij*_ = 1) = *µ*_1_ > *µ*_0_ = 𝔼(*w*_*ij*_|*δ*_*ij*_ = 0). *Let* 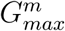 *be the maximum detectable community structure from m simulated graphs, then we will have*

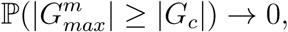

*as n* → ∞.

*Proof*. The claim is automatically true from Theorem 2. □

The above theorem shows that the Type II error of network-level inference is low when phenotype related subnetworks present. Graph combinatorics plays an important role in group-level network statistical inference because it is rare for an organized subnetwork being associated with a phenotype by chance. Given the 𝓁_0_ norm regularization based algorithm can capture the dense subgraphs with a density *γ* ≫ *µ*_0_, the following network-level statistical tests can simultaneously reduce false positive and false negative discovery rates. Besides network-level inference, we can also draw statistical inference on individual edges inside and outside *G*_*c*_ adaptively based on an empirical Bayes framework (Chen et al., 2018). Since this topic is beyond the scope of this article, we include brief descriptions in the supplementary materials. We also prove that the edge-wise false positive and false negative error rates given {*G*_*c*_} are also simultaneously improved compared with the conventional methods that apply a universal cut-off like in FDR, local *fdr*, and FWER.

In summary, GLEN provides multiscale inference (both edge-level and network-level) without prior knowledge of informative subnetworks. The 𝓁_0_ norm regularization can reliably extract latent subnetworks (topological structures) consisting of phenotype-related edges. The detected network topology is naturally linked with our graph combinatorics based inference which is novel, powerful, and unique to group-level network analysis.

## 3 Data example

The primary data set *D*^1^ includes 70 individuals with schizophrenia (age = 40.80 ± 13.63 years) and 70 control subjects (age = 41.79 ± 13.44 years) frequency-matched on age (*t* = 0.62, *p* = 0.54) and sex ratio (*χ*^2^ = 0, *p* = 1). In the validation data set *D*^2^, another 30 individuals with schizophrenia (age = 39.73 ± 13.79 years) and 30 control subjects were recruited (age = 39.73 ± 14.16 years) matched on age (*t* = 0.27, *p* = 0.78) and sex ratio (*χ*^2^ = 0.09, *p* = 0.77), following the initial sample. The recruitment procedures, inclusion and exclusion criteria, and imaging acquisition and preprocessing procedure were kept the same. The details of subject recruitment, imaging acquisition, and preprocessing procedures are described in Supplementary Materials. The nodes of the connectome graph *G* are defined based on the commonly used automated anatomical labeling (AAL) regions. Time courses of all voxels within a 10 mm sphere around the centroid of each region are pre-processed as region-wise signals, followed by calculating 4005 Pearson correlation coefficients between the time courses of the 90 AAL regions. Fishers Z transformation and normalization are then applied to obtain connectivity matrices. We apply GLEN to both data sets separately and then compare *D*^1^ and *D*^2^ results. We also compare the results by GLEN with the traditional edge-wise and the commonly used network based statistic (NBS) methods.

### 3.1 Network-level results of *D*^1^

We first apply GLEN to *D*^1^. Let symmetric matrix **W**^1^ be the whole brain graph edge-wise testing result matrix (Fig 3a), where the element is *w*_*ij*_ = − log(*p*_*ij*_) where i and j are two distinct brain regions and *p*_*ij*_ is the corresponding test p-value for the edge between i and j. The graph combinatorics based testing results show that one subnetwork in *D*^1^ is significant (p *<* 0.001). The significant subnetwork (**R**^1^ denotes the subnetwork from *D*^1^) includes 22 nodes, 231 altered edges, and a well-organized topological structure. The detected topology is a community structure with 22 nodes including the left medial frontal cortex, bilateral insula, bilateral anterior and middle cingulate cortices, bilateral Heschl gyrus and superior temporal cortices, bilateral paracentral and postcentral cortices, right precentral cortex, and the precueous (Fig 3a to 3c) (full list of region names in Table 1 of supplementary materials).

**Table 1:** Region names and coordinates in the subnetwork of *D*^1^

| Region Name | x | y | z |
| --- | --- | --- | --- |
| Precentral R | -39 | -6 | 51 |
| Rolandic Oper L | -47 | -8 | 14 |
| Rolandic Oper R | 53 | -6 | 15 |
| Supp Motor Area L | -5 | 5 | 61 |
| Supp Motor Area R | 9 | 0 | 62 |
| Frontal Sup Medial L | -5 | 49 | 31 |
| Insula L | -35 | 10 | 3 |
| Insula R | 39 | 6 | 2 |
| Cingulum Ant L | -4 | 35 | 14 |
| Cingulum Ant R | 8 | 37 | 16 |
| Cingulum Mid L | -5 | -15 | 42 |
| Cingulum Mid R | 8 | -9 | 40 |
| Postcentral L | -42 | -23 | 49 |
| Postcentral R | 41 | -25 | 53 |
| Precuneus L | -7 | -56 | 48 |
| Paracentral Lobule L | -8 | -25 | 70 |
| Paracentral Lobule R | 7 | -32 | 68 |
| Heschl L | -42 | -19 | 10 |
| Heschl R | 46 | -17 | 10 |
| Temporal Sup L | -53 | -21 | 7 |
| Temporal Sup R | 58 | -22 | 7 |
| Temporal Pole Sup R | 48 | 15 | -17 |

**Figure 2:**
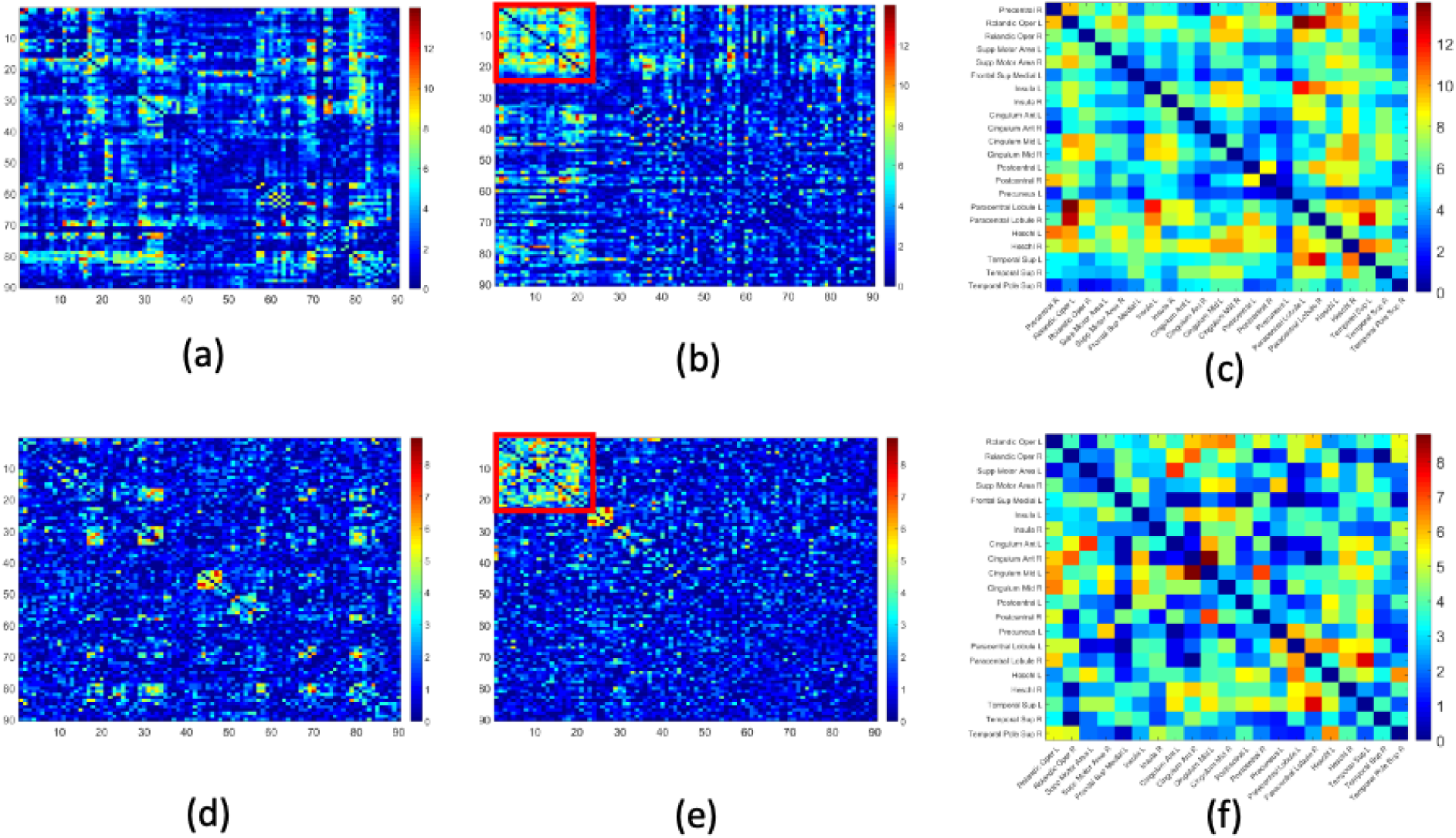
Applying GLEN to clinical data *D*^1^ (a-c) and replication data *D*^2^ (d-f). a) A heatmap of log(*p*) of the first data set (*D*^1^). A hotter pixel indicates more differential edge between cases and controls. There is no apparent topological pattern of these hot edges. b) We then perform GLEN in *D*^1^ and find a significant subnetwork (red square, which is magnified in c). c) The enlarged disease-relevant subnetwork in *D*^1^ with region names. d) A heatmap of log(p) of the second data set (*D*^2^). e) The detected disease-relevant subnetwork by using *D*^2^ alone. f) The enlarged network in *D*^2^ with region names.

**Figure 3:**
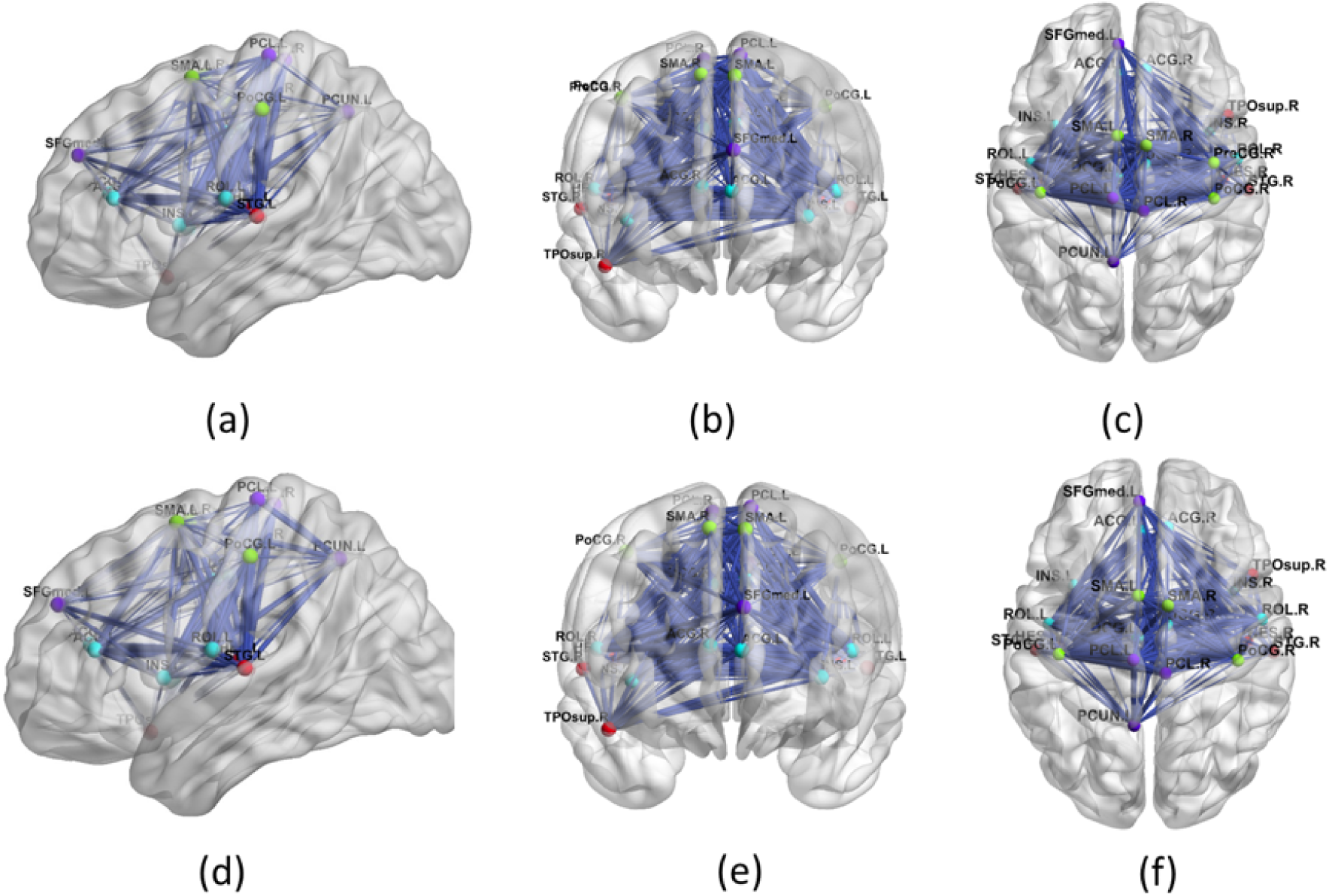
The edges in the subnetwork using 3D demonstration for data set 1 (a)-(c) and data set 2 (d)-(f). The width of length is proportional to the effect size. The disease-relevant network involves with the salience network, part of default mode network, and part of executive network, and more importantly the interconnections between these three networks are revealed. (e)-(f) show 3D brain subnetwork for data set 2, which shows a highly replicable brain subnetwork as seen in data set 1 with one brain region (precentral R) less.

### 3.2 Network-level results of *D*^2^

One subnetwork is significant in *D*^2^ (*p <* 0.001). **R**^2^ (subnetwork from *D*^2^) includes 21 nodes, 210 edges in a clique/community topological structure. The majority of the nodes of **R**^2^ are similar to **R**^1^ with some minor differences, and include left medial superior frontal gyrus, bilateral insula, bilateral anterior and middle cingulate cortices, bilateral Heschl gyrus, Rolandic operculums, supplementary motor areas, paracentral lobules, postcentral lobules, and left precuneus (Fig 3d to 3e) (Table 2 in supplementary materials). Similar to **R**^1^, most (206 of the 210) edges showed reduced connectivity in patients with schizophrenia in **R**^2^.

**Table 2:** Region names and coordinates in the subnetwork of *D*^2^

| Region Name | x | y | z |
| --- | --- | --- | --- |
| Rolandic Oper L | -47 | -8 | 14 |
| Rolandic Oper R | 53 | -6 | 15 |
| Supp Motor Area L | -5 | 5 | 61 |
| Supp Motor Area R | 9 | 0 | 62 |
| Frontal Sup Medial L | -5 | 49 | 31 |
| Insula L | -35 | 10 | 3 |
| Insula R | 39 | 6 | 2 |
| Cingulum Ant L | -4 | 35 | 14 |
| Cingulum Ant R | 8 | 37 | 16 |
| Cingulum Mid L | -5 | -15 | 42 |
| Cingulum Mid R | 8 | -9 | 40 |
| Postcentral L | -42 | -23 | 49 |
| Postcentral R | 41 | -25 | 53 |
| Precuneus L | -7 | -56 | 48 |
| Paracentral Lobule L | -8 | -25 | 70 |
| Paracentral Lobule R | 7 | -32 | 68 |
| Heschl L | -42 | -19 | 10 |
| Heschl R | 46 | -17 | 10 |
| Temporal Sup L | -53 | -21 | 7 |
| Temporal Sup R | 58 | -22 | 7 |
| Temporal Pole Sup R | 48 | 15 | -17 |

### 3.3 Comparing subnetworks in *D*^1^ and *D*^2^

Remarkably, we note that **R**^2^ ⊂ **R**^1^: GLEN reports the altered subnetwork in *D*^1^ that can be rediscovered when analyzing *D*^2^ independently with only one node in difference. Based on the combinatorics, ℙ(**R**^2^ ⊂ **R**^1^) *<* 2×10^−16^ given 21 nodes are included in **R**^2^. Although the sample size of *D*^2^ is smaller and the testing p-values are larger, the subnetwork detection in *D*^2^ is not affected by the sample size and other sources of noise. While the statistical inference on individual edges is subject to numerous variations and false positive-negative findings, we found that the latent network topological patterns of differentially expressed edges are stable across independent samples. The new network size regularization term is critical to recognize the organized patterns from the noisy background. The organized (latent) subnetwork topological structure can reduce the false positive and false negative findings simultaneously. As a result, the subnetwork level findings are reproducible and verified by *D*^2^.

### 3.4 Comparisons with existing methods

For comparison purpose, we also perform edge-wise multiple testing analysis and NBS. Wilcoxon rank sum tests are then used to assess patient-control differences in the normalized correlation coefficients for all edges. In *D*^1^, 430 of 4005 edges are *p <* 0.005. *p* = 0.005 is commonly used for uncorrected threshold in neuroimaging literature (Derado biometrics 2010). After FDR correction for multiple comparison, none of 4005 edges is found significant by using the threshold *q* = 0.05. In *D*^2^, 22 of 4005 edges are *p <* 0.005, and none of edges are found significant after applying FDR correction with the threshold *q* = 0.05. Two edges among the 430 edges in *D*^1^ and 22 edges in *D*^2^ overlap, which indicates a very low replicability between the two data sets. In addition, the conventional network method NBS shows no differentially expressed subnetwork in *D*^1^ and *D*^2^ by using various thresholds (tuning threshold values from 2 to 6). This is likely due to the loss of power by the impact of false positive edges (without 𝓁_0_ norm shrinkage).

Finally, the positive agreement is used to compare the reproducibility of features between *D*^1^ and *D*^2^ using GLEN vs. individual edge based statistics. The network approach is significantly better than the individual edge based method. In summary, by utilizing an independent replication data set collected posteriorly we can conclude that the findings identified by the GLEN are more reproducible.

### 3.5 Biological interpretation of the brain subnetwork

The brain region constellation of the detected subnetwork consists of many well-known brain regions involved with the schizophrenic disorder, which included inferior frontal, superior temporal, insula, cingulate, and paracentral areas (Fig 3). This altered subnetwork is composed of approximately the salience network (SN), part of default mode network (DMN), and part of central executive network (CEN), which have been frequently associated with abnormalities in schizophrenia during attention, working memory, and executive control, and resting functional imaging studies. Interestingly, the detected subnetwork reveals not only that the SN, DMN, and CEN are altered but also that the interconnections between these three networks are disrupted. Of the 231 differentially expressed edges, all edges show decreased or equivalent connections in patients with schizophrenia. This aligns with findings suggesting that schizophrenia is a ‘dysconnectivity’ disorder with primarily reduced functional connectivity across brain regions (Lynall et al., 2010), although medication effects cannot be completely ruled out.

## 4 Simulations

In the simulation analysis, we investigate whether graph combinatorics based statistical inference can reduce false positive and false negative error rates for multivariate edge analysis. We simulate group-level connectome data sets ***A*** = {***A***^1^, …, ***A***^*S*^} and the corresponding graph whole brain connectome *G*, we define a community subnetwork *G*_*c*_ ⊂ *G* where edges are differentially expressed between controls and cases. We let |*V* | = 100, |*V*_*c*_| = 20, and *G*_*c*_ to be a clique (and |*E*_*c*_| = 190). The transformed (e.g. Fishers Z) connectivity metric of each edge is set to marginally follow a normal distribution with *µ*_0_ and 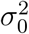 (for controls) and *µ*_1_ and 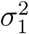 (for cases). For differentially expressed edges (i.e. *e*_*i,j*_ ∈ *E*_*c*_), *µ*_0_ = *µ*_1_ +*θ* and otherwise *µ*_0_ = *µ*_1_. Also, we let 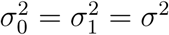. *θ* = 0.5 is used with varying values of *σ*^2^ for different signal-to-noise ratios (SNRs)/standardized effect sizes. By shuffling the order of nodes in *G*, the altered connectivity subnetwork becomes latent in the sample data. Two sample sizes (60 and 120) are used to represent the commonly observed sample size from a single study. Each setting is simulated with different *θ, σ* and the number of subjects for each group for 100 times.

The subnetwork detection algorithm and graph combinatorics based tests of GLEN are applied to each simulated data set. The permutation tests are performed by shuffling the group labels and edge orders to simulate the null, denoted by GLEN1 and GLEN2 (see details in the supplementary materials). The false positive discovery rates (FP) and false negative discovery rates (FN) are evaluated at both subnetwork and edge level rates. Note that edge-wise power can be further calculated as 1 − *FN*. Our method is compared with other multiple testing methods including BenjaminiHochberg FDR and local false discovery rate control (*fdr*). The false positive findings (number of FP edges in mean and standard deviation across 100 repetitions) and false negative (FN) edges are shown in Table 1.

GLEN shows improved performance on network level and edge level inference by identifying the latent and differentially expressed subnetwork with 0 FP and FN rates. Next, GLEN (based on the selected subnetwork) is compared to FDR and local fdr at individual edge inference using *q* = 0.2 as the cut-off for both FDR and fdr. The results show that generally FDR has higher FP but lower FN rates compared with fdr (i.e. fdr is more conservative). Importantly, GLEN outperforms FDR and fdr when jointly considering FP and FN rates, see Table 1. Finally, we apply NBS method to the simulated data, but no subnetwork is detected by NBS thus the power is 0 when tuning the cutoff parameter from 3 to 6 for all settings (not shown in Table 1). One possible reason is that the breadth first algorithm in NBS can include many false positive edges and thus lose power in the permutation test.

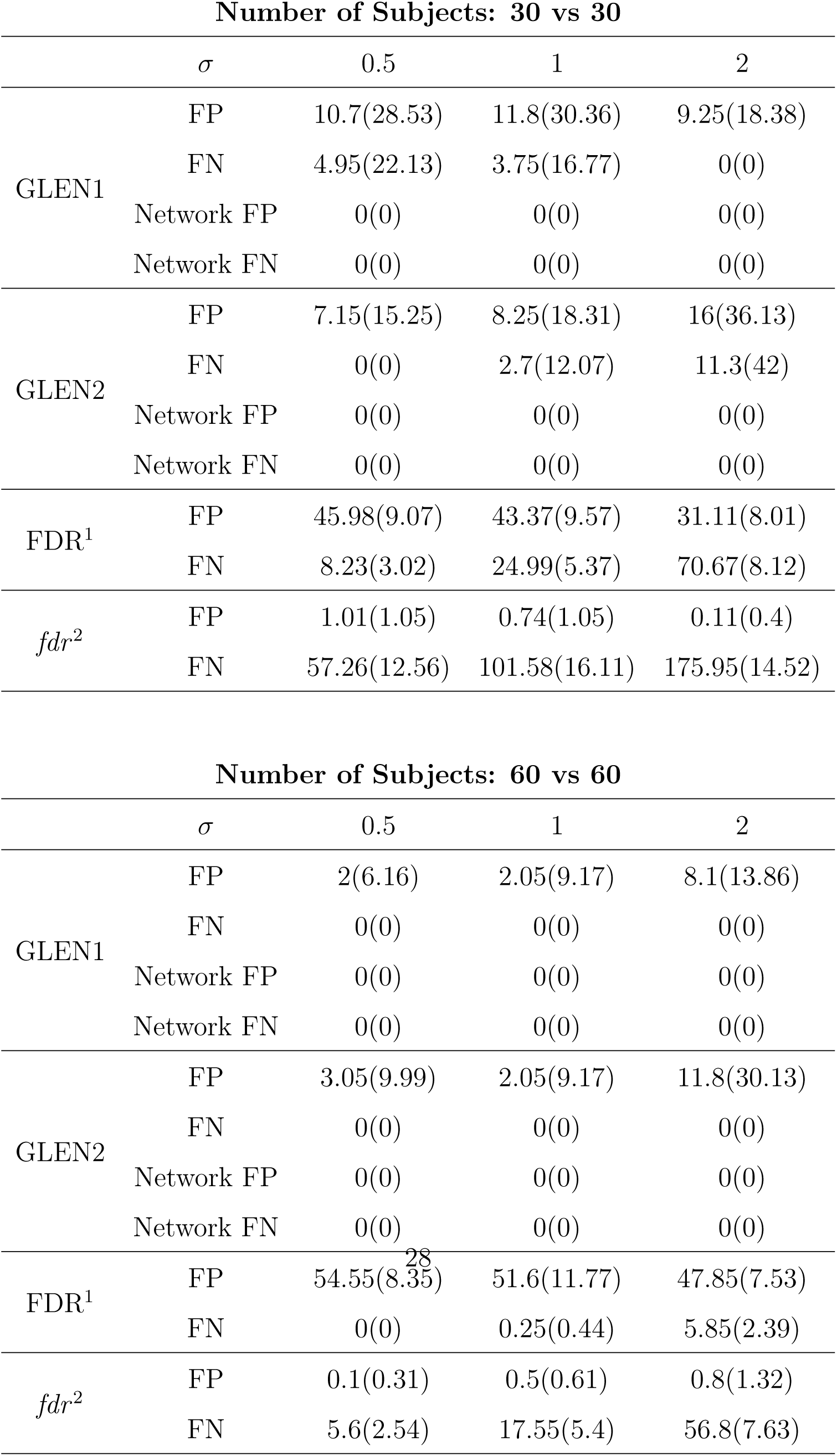

On Type I error rate, we count the number of false positive significant subnetworks for the data sets with no differentially expressed connectome networks (e.g. *θ* = 0). Based on simulation of 1000 iterations, the network level false positive rate of GLEN1 is 1.2% and GLEN2 is 2.9%. Therefore, the network level Type I error is well controlled and below the subnetwork claimed level of 5%.

## 5 Discussion

In this article, we present a graph combinatorics based approach for group-level network inference which is motivated by brain connectome research. The importance of multiple testing problem in neuroimaging data cannot be overstated, due to the urgent need to improve the reliability and reproducibility of findings in neuroscience research (Lazar, 2008; Lindquist, 2008; Eklund et al., 2016). GLEN capitalizes on latent organized network topology of multivariate edge variables and performs graph combinatorics based tests yielding network and edge level inference results with increased power and reduced false positive discovery rates.

We develop a new objective function to extract latent subnetworks via 𝓁_0_ norm shrinkage that is specifically tailored for a noisy and less dense input matrix **W**. 𝓁_0_ norm shrinkage for edge variables in a graph space is fundamentally different from the conventional settings of high-dimensional data analysis (e.g. LASSO), which reflects the difference between the 𝓁_0_ norm of a vector and a subgraph. We further report that our 𝓁_0_ norm regularization can be implemented efficiently. The 𝓁_0_ norm regularization ensures that detected subnetworks are dense subgraphs where informative edges are highly concentrated in organized topological structures. The combinatorial probability of an organized topological structure in a random graph model is essentially zero. Thus, the graph combinatorics based tests can be utilized to examine the hypothesis whether networks are systematically influenced by the phenotype. The graph combinatorics based method is particularly useful for group-level network inference because it produces both edge and network level inference and reveals biologically meaningful network topology.

GLEN may also provide a solution to improve the reproducibility and replicability of findings across studies in neuroscience research (Eklund et al., 2016). One potential reason for the low replicability is the universal threshold of multiple testing correction methods (e.g. the primary thresholding and FDR/FWER methods). When noise and heterogeneity present, the universal threshold can either have a low false discovery rate or high power/sensitivity, but not both. Therefore, the results for each study either include a small proportion of true signals being discovered (low sensitivity) or a small proportion of discoveries that are true positive signals (a low true discovery rate), and the chance for the true discoveries overlap with each other across studies is very rare i.e. low replicability. Our results demonstrate that findings by GLEN are highly replicable because both false positive and false negative findings are simultaneously reduced via graph combinatorics based statistical inference. The 𝓁_0_ shrinkage algorithm is designed to detect topological structures of true positive signals with very small graph combinatorial probability. The detected topological structures, in turn, can be explicitly used in graph combinatorics based statistical inference. In that, edge variables do not only borrow strength from each other but also they consolidate into a unified topological structure to achieve much improved inferential accuracy. In summary, we develop a graph-combinatorics based group-level network analysis method GLEN which can yield accurate multivariate inference and provide novel insights of network topology.

## SUPPLEMENTARY MATERIALS

### A1. Testing the nonrandom patterns of differentially expressed edges

Let *r*_0_ be a threshold for multiple testing correction, and *E*_1_(*r*_0_) = {*e*_*i,j*_| |*w*_*ij*_| > *r*_0_} and the corresponding edge-induced subgraph *G*_1_(*r*_0_) ⊂ *G. d*_*i*_ = Σ_*j* ≠ *i*_ *I*(|*w*_*ij*_| > *r*_0_) represents the degree of node *i* in *G*_1_. We examine *H*_0_ : *G*_1_ is a random graph vs. *H*_1_ : *G*_1_ is not random. However, the set of edges with {***β***_*ij*_ ≠ 0} is unknown and thus needs to be estimated. Under *H*_0_, the multiple correction based thresholding is valid because *h*_*ij*_ = *h*_*i′j′*_ |*w*_*ij*_ = *w*_*i′j′*_ and the inference is irrelevant to the shuffling order of edges.

Under *H*_0_ that *G*_1_ is a random graph, *d*_*i*_ follows a Poisson distribution (Newman 2002). Thus, we reject *H*_0_ if the distribution *d*_*i*_ statistical significantly deviate from the Poisson distribution. We perform permutation test to assess the significance of the deviation. Since the true *r*_0_ is unknown, we assume it follows a distribution *f* (*r*_0_). *P* (*d*_*i*_) denotes the empirical sample distribution of *d*_*i*_ and *Q*(*d*_*i*_) = Poisson(*d*_*i*_) with parameter estimated by the random graph model. The Kullback-Leibler divergence based statistic 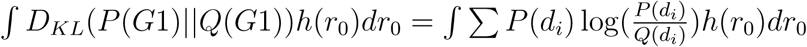 is used to measure the deviation. *h*(*r*_0_) is the prior distribution of the cutoff. In each permutation, *G*_1_(*r*_0_) is randomized by shuffling the order of edge is shuffled (Hanhijarvi 2009). We reject the null if the the observed testing statistic is among the top 5 percentile of statistics from permutations. If we fail to reject *H*_0_, the conventional methods like FDR can be used. If *H*_0_ is rejected, our next goal is to identify the and test latent topological structure of *G*_1_

### A2. Derivation of the algorithm for objective function (1)

The optimization of objective function (1) is implemented by exhaustive search for *C* and estimating **Û** at each *C*. With a given *C*,

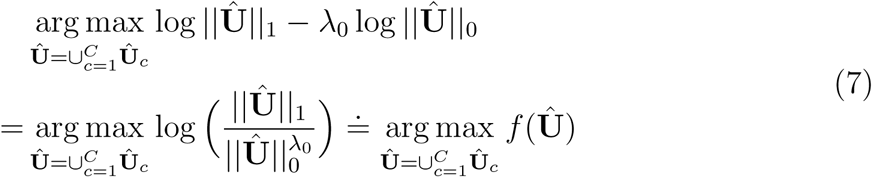

By default *λ*_0_ = 0.5 reflects balanced covering quality and quantity of true positive edges, and the objective function 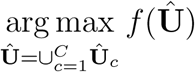 then becomes the well-known problem of *k* dense subgraph discovery, where *f* (·) is the density function. The problem has been solved in polynomial time by Goldberg’s min-cut algorithm (Goldberg, 1984) and a greedy algorithm with 1/2 approximation by Charikar (2000). In addition, the default topological community structure can be considered as quasi-cliques and the problem can be solved by additive approximation algorithms and local-search heuristics (Tsourakakis et al., 2013). Alternatively, with the mild spatially-invariant assumptions that 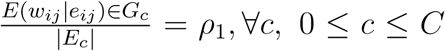, and 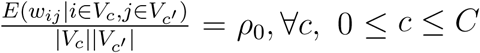 the primary objective function is equivalent to

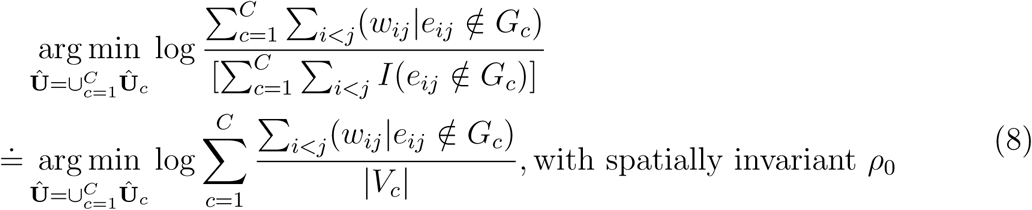

Although the objective function (8) is not convex, the issue of local optima in the discrete optimization can be solved by restarting the algorithm for several times with different initializations and/or through orthonormal transforms (Stella and Shi, 2003 and Bolla, 2013). The proposed algorithm may better extract multiple weighted dense subgraphs (with an unknown number and unknown sizes of dense subgraphs) than the existing algorithms of dense subgraph discovery (Chen et al., 2018). We then choose the optimal *C*\* by grid searching that maximizes the following criteria:

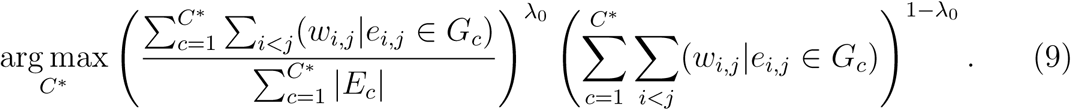

The criteria (9) can be directly derived from the our primary objective function that 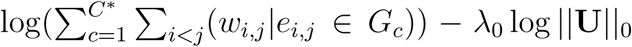. The first term in (9) indicates the ‘qaultiy’ (the area density) of the extracted subgraphs, while the second term represents the ‘quantity’ of edges covered by the subgraphs. *C*\* is selected with optimal quality and quantity in terms of covering informative edges. *λ*_0_ can be tuned to either extract subgrahs with higher area density (i.e. low false positive rates) or covering more high-weight edges using subgraphs with larger sizes (i.e. low false negative rates). In general, *C*\* selection is robust for *λ*_0_ in the range of 0.4 to 0.7.

In summary, the above procedure can extract latent organized topological structures containing most high-weight edges while controlling the sizes of the topological structures by 𝓁_0_ norm regularization. Furthermore, we have recently developed more flexible algorithms to extract subgraphs beyond the default community structure, for example, k-partite/rich club and interconnected induced subugrahs can be further detected based on detected quasi-cliques (Chen et al., 2019 and Wu 2019). These more sophisticated topological structures can further improve the objective function by preserving the high-weight edges inside of more parsimonioulsy-sized subgraphs. The sample codes can be found at https://github.com/shuochenstats/Network_program/tree/master/GLEN_package.

### A3. Proof of Lemma 2

Based on the derivation of algorithm in Appendix A2, it suffices to show the consistency results are guaranteed for spectral clustering in our setting of a continuous stochastic block model. The proof of theorem 3.1 in Lei et al. (2015) can be easily extended to a weighted case using continuous versions of Bernstein inequality and Chernoff bounds.

To bound light pairs, 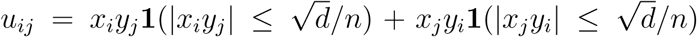, then 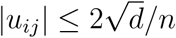, and *x*^*T*^ *W* ′*y* can be written as

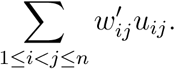

Then, for zero-mean independent random variables, apply Bernstein inequality,

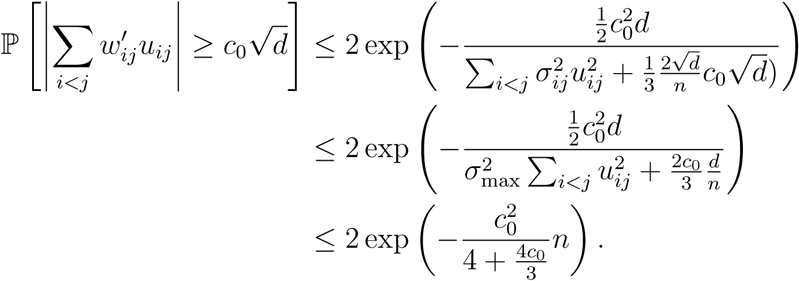

In bounding heavy pairs, let *e*(*I, J*) be the summation of edge weights in node sets I and J: 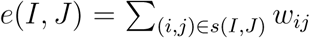. Define 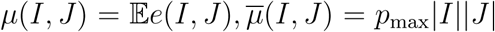. We could obtain continuous versions of Lemma 4.1 and 4.2 in supplementary material of Lei et al. (2015).

Using Bernstein inequality:

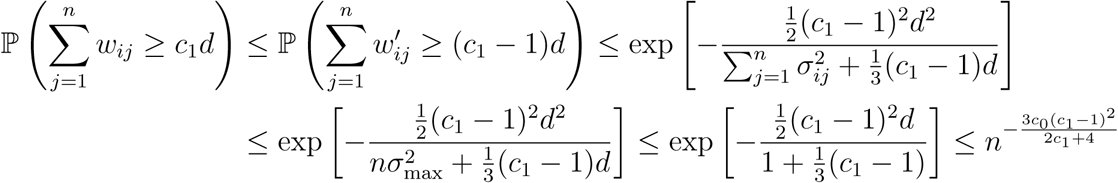

We have for *c*_0_ > 0, there exists constant *c*_1_ = *c*_1_(*c*_0_) such that with probability at least 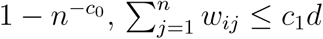.

From Chernoff Bound:

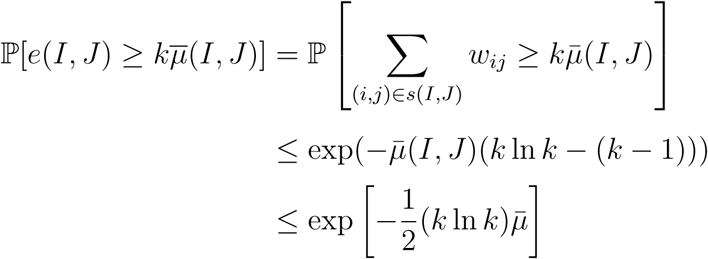

the lemma 4.2 is true from exacly the same calculations.

Hence, our claim is true with stated assumptions from theorem 3.1 of Lei et al. (2015). □

### A4. Proof of Theorem 1

*Part 1* We first consider the case *C* = *C*\* − 1. Then, intuitively, the estimated labels 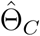 should be obtained by merging two smallest communities under Θ*. Without loss of generality, we assume it merges the last two communities with label *C*\* − 1 and *C*\*. The associated subgraph-based matrices changed from ***Û***^*\**^ to ***Û***^*C*^. Denote *X*_*i*_ ∼ *f*_1_, and *Y*_*i*_ ∼ *f*_0_ denote random variables for phenotype-related and unrelated edges. Therefore, from lemma 1,

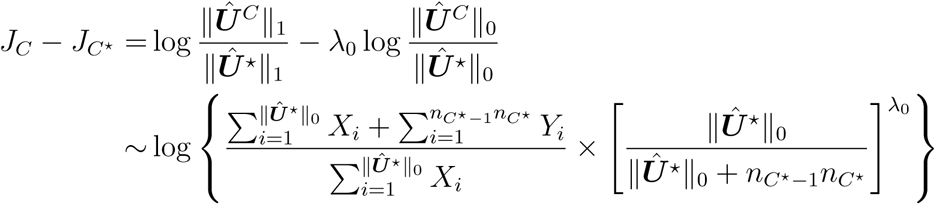

Then, from chernoff bounds, for any *δ* > 0

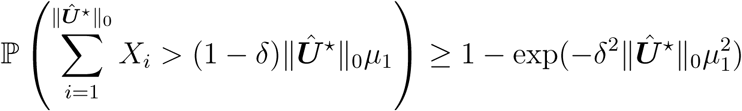

and

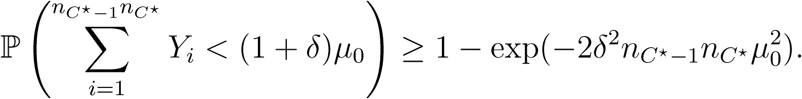

Hence,

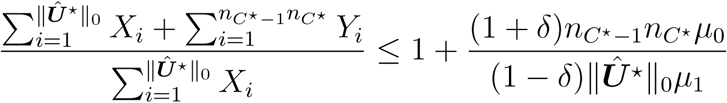

with probability greater than

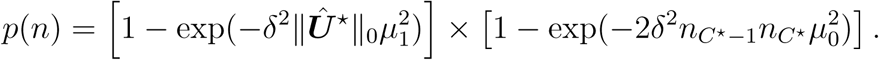

Thus, with probability *p*(*n*),

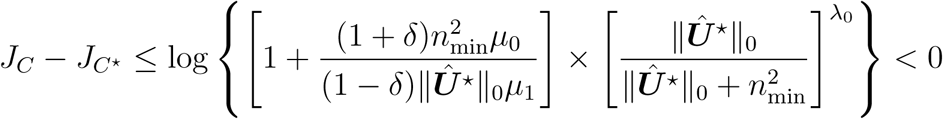

as *n* → ∞, with *λ*_0_ = 1*/*2 and

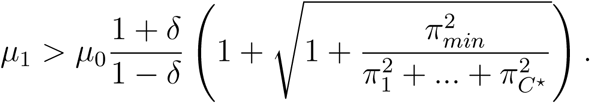

Inductively, we could conclude for *C < C*\*, ℙ(*J*_*C*_ *< J*_*C*_*\**) → 1.

*Part 2* For cases *C* > *C*\*, assume for each subgraph *V*_*c*_, *c* = 1, …, *C*\*, *Ŝ*_*c*_ is the corresponding estimated nodes set with smallest assignment error:

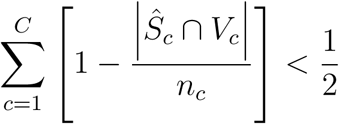

among all permutations. Then, the least favorable case (the case with highest *J*_*C*_) happens when all mis-assignment nodes should belong to the smallest community (*n*_min_). Thus, let *n*_*e*_ denote the number of mis-assigned nodes for the smallest community, we have

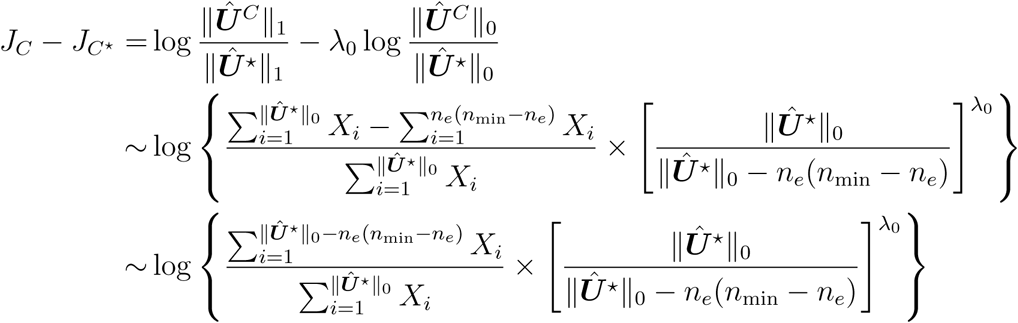

When *n*_*e*_ = *O*_*p*_(*n*_min_), from multinomial distribution of subgraph sizes, we have 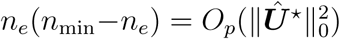, and ℙ(*J*_*C*_ *< J*_*C*_*\**) → 1 as *n* → ∞. Hence, for grid searched 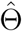, the mis-assignment nodes (up to a permutation) should satisfy *n*_*e*_ = *o*_*p*_(*n*_min_). The case with

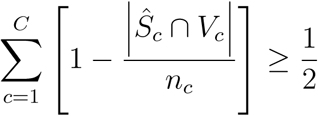

has equivalent least favorable case considering permutations. □

### A5. Lemma 3

For a graph *G*(*V, E*), a subgraph indexed by *S* ⊆ *V* is said to be a *γ*−quasiclique, if the subgraph has at least 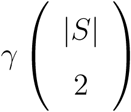 edges, where *γ* ∈ [0, 1] is a parameter. For a random graph *G*(*n, p*) and 1 > *γ* > *p* > 0, let 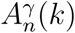 be the maximum number of edges that can be included in at most *k γ*−quasi-clique. Then, the maximum number of edges that can be included in at most ln *n γ*−quasi-clique is 𝒪((ln *n*)^3^), in other words,

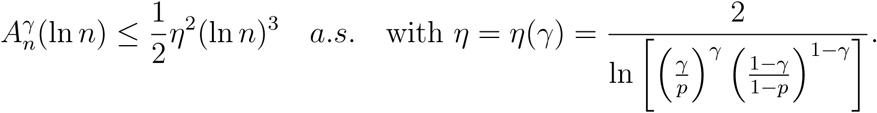

**Proof**. If the number of edges included in at most *k γ*−quasi-clique is 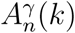, there must be at least one *γ*−quasi-clique which include edges no less than 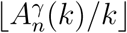. From Theorem 1 in Veremyev et al. (2012),

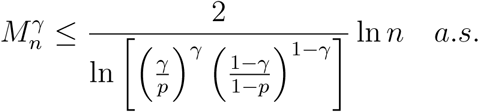

where 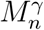 is the maximum number of vertices in a *γ*−quasi-clique. Hence,

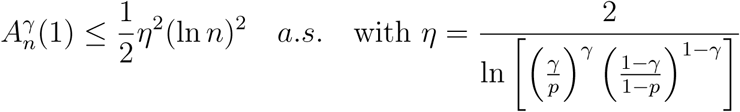

Therefore, 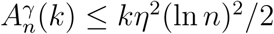 almost surely.

### A6. Edge-level inference by GLEN

GLEN provides network level inference by reporting phenotype related subnetwork *G*_*c*_ instead of *e*_*ij*_ ∈ *G*_*c*_. We argue that the detected subnetworks can also assist edge-level inference by applying an empirical Bayes based adaptive thresholding strategy. The details of method is introduced in section 2.2 Chen et al. (2018). When phenotype related edges consist of organized topological structures, the false positive and false negative discovery rates for individual edges are lower than conventional universal threshold multiple testing methods.

#### Theorem 4

Let *F*_0_(*x*) = *P* (*w*_*ij*_ ≤ *x*|*δ*_*ij*_ = 0) and *F*_1_(*x*) = *P* (*w*_*ij*_ ≤ *x*|*δ*_*ij*_ ≠ 0). Under the conditions,

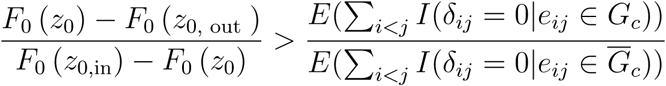

and

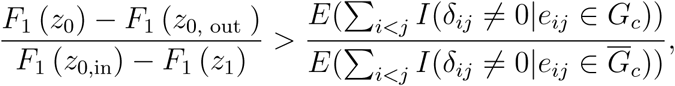

where *z*_0_ is a universal threshold value and *z*_0,in_, *z*_0,out_ are threshold values inside and outside the community structure, we have

1. the expected false positively thresholded edges by using the network level inference (GLEN) are less than the universal thresholding method

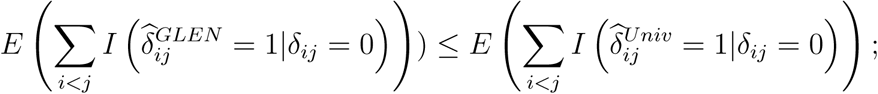
2. the expected false negatively thresholded edges by using GLEN are less than the universal thresholding method

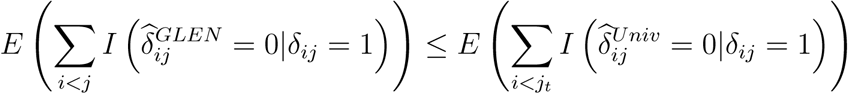

*Proof*. See the proof of Theorem 1 in Chen et al. (2018). □

### A7. Permutation test

#### Graph edge permutation vs. graph node permutation

There are two sets of elements in a graph: the set of vertices and the set of edges as in *G* = {*V, E*}. Correspondingly, there are two options of permutation: permuting nodes or edges. First, we consider the permutation of nodes as a reordering process*π*, which is an ‘edge-preserving bijection. If two nodes *a* and *b* are connected in graph*G*, then in the node-permuted graph*H* = *π*(*G*) = {*π*(*V*), *F*}, then *π*(*a*) and *π*(*b*) are connected:*E*_*ab*_ = 1 ⇔ *F*_*π*(*a*)*π*(*b*)_ = 1. *G* and H are isomorphic graphs *G* ∼ *H* (see Figs 1d and 1e). The GLEN subnetwork detection algorithms reorder and group the nodes to uncover these latent topological patterns. In contrast, the edge permutation is different because it permutes the order of edges and is not ‘edge-preserving bijection. For example, two nodes *a* and *b* are connected in a graph*G*; the edge-permuted graph *L*(e.g. by permutation*µ*) that *L* = *µ*(*G*) = {*V, F* = *µ*(*E*)} and in *L,a* and *b* are only connected with probability of*p*_*G*_, where *p*_*G*_ is number of connected edges in *G* divided by*n* × *n/*2. Therefore, the above two events are independent: {*E*_*ab*_ = 1}⊥{*F*_*ab*_ = 1}. Hence, though there is an organized pattern in*G*, the edge permuted graph *L* = *µ*(*G*) becomes a random graph without any organized patterns. The connectivity testing p-value matrix after edge permutation represents a random graph where each edge has the identical probability such that*p*_*i,j*_ *< p*_0_. Therefore, the edge permutation can be used to test the organized topological pattern. The GLEN1 permutation test simulates the null by shuffling the group labels (i.e. the order of the covariate of interest) while GLEN2 permute the order of edge to simulate a random graph.

#### Network level test statistic

We propose a new test statistic in the permutation tests, which is specifically tailored for the network as an object. Our goal is to select disease-related subnetwork that is both informative (including the largest number of informative edges that is possible) and efficient (most edges in each subnetwork are significant). Generally, these two factors are conflicting: including more informative edges (resulting in larger networks) can reduce the average significance of edges in the subnetwork whereas restricting only very significant edges in the subnetwork can shrink the network size and overall include less informative edges in *G*. We develop a new test statistic to integrate these two aspects of the network object. The new test statistic

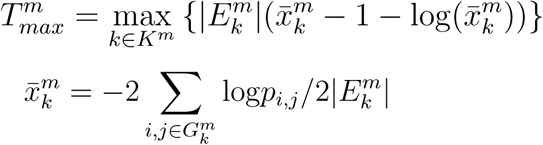

is derived based on Fishers combination test and Chernoff bound of *χ*^2^ the cumulative distribution function. The test statistic is negative (and generally the network is not significant) when it includes a small proportion of informative edges. Therefore, the test statistic jointly evaluates the effect size of edges in the subnetwork and the size of each selected subnetwork which together are an index of the information of a subnetwork.

### A8. fMRI data acquisition and pre-processing

All participants provided written informed consent that had been approved by the University of Maryland Internal Review Board. All participants were evaluated using the Structured Clinical Interview for the DSM-IV diagnoses. We recruited medicated patients with an Axis I diagnosis of schizophrenia through the Maryland Psychiatric Research Center and neighboring mental-health clinics. We recruited control subjects, who did not have an Axis I psychiatric diagnosis, through media advertisements. Exclusion criteria included hypertension, hyperlipidemia, type 2 diabetes, heart disorders, and major neurological events, such as stroke or transient ischemic attack. Illicit substance and alcohol abuse and dependence were exclusion criteria. Data were acquired using a 3-T Siemens Trio scanner equipped with a 32 channel head coil at the University of Maryland Center for Brain Imaging Research. A T1-weighted structural image (MP-RAGE: 1 mm isotropic voxels, 256 × 256 mm FOV, TR/TE/TI = 1900/3.45/900ms) was acquired for anatomical reference. Fifteen minutes of rfMRI was collected on each subject. During the resting scans, subjects were given a simple instruction to rest and keep their eyes closed. Head motion was minimized using foam padding, foam molding, and tapes. RfMRI were acquired over 39 axial, interleaving slices using a gradient-echo EPI sequence (450 volumes, TE/TR = 27/2000 ms; flip angle = 90^*o*^; FOV = 220×220 mm; image matrix = 128×128; in-plane resolution 1.72×1.72mm. Following the previously published procedures, data were preprocessed in AFNI and MATLAB (MathWorks, Inc., Natick, MA). Volumes were slice-timing aligned and motion corrected to the base volume that minimally deviated from other volumes using an AFNI built-in algorithm. After linear detrending of the time course of each voxel, volumes were spatially normalized and resampled to Talairach space at 3×3×3 mm3, spatially smoothed (FWHM 6 mm), and temporally low-pass filtered (fcutoff = 0.1 Hz). For functional connectivity analyses, the six rigid head-motion parameter time courses and the average time course in white matter were treated as nuisance covariates. A white matter mask was generated by segmenting the high-resolution anatomical images and down-gridding the obtained white matter masks to the same resolution as the functional data. These nuisance covariates regress out fluctuations unlikely to be relevant to neuronal activity.

### A9. Tables of brain regions

In the following tables, we list the region names and coordinates of subnetworks from *D*^1^ and *D*^2^.

## Footnotes

1 FDR: Benjamini-Hochberg false discovery rate control (FDR)

2 *fdr* : local false discovery rate control.

## References

Benjamini, Y. and Y. Hochberg (1995). Controlling the false discovery rate: a practical and powerful approach to multiple testing. Journal of the Royal statistical society: series B (Methodological) 57 (1), 289–300.

Bolla, M. (2013). Spectral clustering and biclustering: Learning large graphs and contingency tables. John Wiley & Sons.

Bowman, F. D. (2005). Spatio-temporal modeling of localized brain activity. Biostatistics 6 (4), 558–575.

Cai, T., H. Li, J. Ma, and Y. Xia (2018). Differential markov random field analysis with an application to detecting differential microbial community networks. Biometrika.

Cao, X., B. Sandstede, and X. Luo (2019). A functional data method for causal dynamic network modeling of task-related fmri. Frontiers in neuroscience 13.

Charikar, M. (2000). Greedy approximation algorithms for finding dense components in a graph. In International Workshop on Approximation Algorithms for Combinatorial Optimization, pp. 84–95. Springer.

Chen, S., F. D. Bowman, and Y. Xing (2019). Detecting and testing altered brain connectivity networks with k-partite network topology. Computational Statistics & Data Analysis.

Chen, S., J. Kang, Y. Xing, and G. Wang (2015). A parsimonious statistical method to detect groupwise differentially expressed functional connectivity networks. Human brain mapping 36 (12), 5196–5206.

Chen, S., J. Kang, Y. Xing, Y. Zhao, and D. K. Milton (2018). Estimating large covariance matrix with network topology for high-dimensional biomedical data. Computational Statistics & Data Analysis 127, 82–95.

Chen, S., Y. Xing, J. Kang, P. Kochunov, and L. E. Hong (2018). Bayesian modeling of dependence in brain connectivity data. Biostatistics.

Craddock, R. C., S. Jbabdi, C.-G. Yan, J. T. Vogelstein, F. X. Castellanos, A. Di Martino, C. Kelly, K. Heberlein, S. Colcombe, and M. P. Milham (2013). Imaging human connectomes at the macroscale. Nature methods 10 (6), 524.

Derado, G., F. D. Bowman, and C. D. Kilts (2010). Modeling the spatial and temporal dependence in fmri data. Biometrics 66 (3), 949–957.

Durante, D., D. B. Dunson, et al. (2018). Bayesian inference and testing of group differences in brain networks. Bayesian Analysis 13 (1), 29–58.

Efron, B., T. Hastie, I. Johnstone, R. Tibshirani, et al. (2004). Least angle regression. The Annals of statistics 32 (2), 407–499.

Eklund, A., T. E. Nichols, and H. Knutsson (2016). Cluster failure: Why fmri inferences for spatial extent have inflated false-positive rates. Proceedings of the national academy of sciences 113 (28), 7900–7905.

Fan, J. and J. Lv (2008). Sure independence screening for ultrahigh dimensional feature space. Journal of the Royal Statistical Society: Series B (Statistical Methodology) 70 (5), 849–911.

Ginestet, C. E., J. Li, P. Balachandran, S. Rosenberg, E. D. Kolaczyk, et al. (2017). Hypothesis testing for network data in functional neuroimaging. The Annals of Applied Statistics 11 (2), 725–750.

Gionis, A. and C. E. Tsourakakis (2015). Dense subgraph discovery: Kdd 2015 tutorial. In Proceedings of the 21th ACM SIGKDD International Conference on Knowledge Discovery and Data Mining, pp. 2313–2314. ACM.

Goldberg, A. V. (1984). Finding a maximum density subgraph. University of California Berkeley, CA.

Hazimeh, H. and R. Mazumder (2018). Fast best subset selection: Coordinate descent and local combinatorial optimization algorithms. arXiv preprint 1803.01454.

Higgins, I. A., Y. Guo, S. Kundu, K. S. Choi, and H. Mayberg (2018). A differential degree test for comparing brain networks. arXiv preprint 1809.11098.

Kim, Y. (2014). Convolutional neural networks for sentence classification. arXiv preprint 1408.5882.

Kundu, S., J. Ming, J. Pierce, J. McDowell, and Y. Guo (2018). Estimating dynamic brain functional networks using multi-subject fmri data. NeuroImage 183, 635–649.

Lazar, N. (2008). The statistical analysis of functional MRI data. Springer Science & Business Media.

Lei, J., A. Rinaldo, et al. (2015). Consistency of spectral clustering in stochastic block models. The Annals of Statistics 43 (1), 215–237.

Li, R., W. Zhong, and L. Zhu (2012). Feature screening via distance correlation learning. Journal of the American Statistical Association 107 (499), 1129–1139.

Li, X., S. Xie, D. Zeng, and Y. Wang (2018). Efficient 0-norm feature selection based on augmented and penalized minimization. Statistics in medicine 37 (3), 473–486.

Lindquist, M. A. (2008). The statistical analysis of fmri data. Statistical science 23 (4), 439–464.

Lukemire, J., S. Kundu, G. Pagnoni, and Y. Guo (2017). Bayesian joint modeling of multiple brain functional networks. arXiv preprint 1708.02123.

Lynall, M.-E., D. S. Bassett, R. Kerwin, P. J. McKenna, M. Kitzbichler, U. Muller, and E. Bullmore (2010). Functional connectivity and brain networks in schizophrenia. Journal of Neuroscience 30 (28), 9477–9487.

Miyauchi, A. and N. Kakimura (2018). Finding a dense subgraph with sparse cut. In Proceedings of the 27th ACM International Conference on Information and Knowledge Management, pp. 547–556. ACM.

Narayan, M., G. I. Allen, and S. Tomson (2015). Two sample inference for populations of graphical models with applications to functional connectivity. arXiv preprint 1502.03853.

Newman, M. E. and M. Girvan (2004). Finding and evaluating community structure in networks. Physical review E 69 (2), 026113.

Risk, B. B., M. C. Kociuba, and D. B. Rowe (2018). Impacts of simultaneous multislice acquisition on sensitivity and specificity in fmri. NeuroImage 172, 538–553.

Rohe, K., S. Chatterjee, B. Yu, et al. (2011). Spectral clustering and the high-dimensional stochastic blockmodel. The Annals of Statistics 39 (4), 1878–1915.

Shaddox, E., C. B. Peterson, F. C. Stingo, N. A. Hanania, C. Cruickshank-Quinn, K. Kechris, R. Bowler, and M. Vannucci (2018). Bayesian inference of networks across multiple sample groups and data types. Biostatistics.

Shen, X., W. Pan, and Y. Zhu (2012). Likelihood-based selection and sharp parameter estimation. Journal of the American Statistical Association 107 (497), 223–232.

Simpson, S. L., M. Bahrami, and P. J. Laurienti (2019). A mixed-modeling framework for analyzing multitask whole-brain network data. Network Neuroscience 3 (2), 307–324.

Stella, X. Y. and J. Shi (2003). Multiclass spectral clustering. In null, pp. 313. IEEE.

Tsourakakis, C., F. Bonchi, A. Gionis, F. Gullo, and M. Tsiarli (2013). Denser than the densest subgraph: extracting optimal quasi-cliques with quality guarantees. In Proceedings of the 19th ACM SIGKDD international conference on Knowledge discovery and data mining, pp. 104–112. ACM.

Veremyev, A., V. Boginski, P. A. Krokhmal, and D. E. Jeffcoat (2012, 12). Dense percolation in large-scale mean-field random networks is provably explosive. PLOS ONE 7 (12), 1–14.

Wang, W., X. Zhang, and L. Li (2019). Common reducing subspace model and network alternation analysis. Biometrics.

Warnick, R., M. Guindani, E. Erhardt, E. Allen, V. Calhoun, and M. Vannucci (2018). A bayesian approach for estimating dynamic functional network connectivity in fmri data. Journal of the American Statistical Association 113 (521), 134–151.

Xia, C. H., Z. Ma, Z. Cui, D. Bzdok, D. S. Bassett, T. D. Satterthwaite, R. T. Shinohara, and D. M. Witten (2019). Multi-scale network regression for brain-phenotype associations. bioRxiv, 628651.

Xia, Y. and L. Li (2017). Hypothesis testing of matrix graph model with application to brain connectivity analysis. Biometrics 73 (3), 780–791.

Xia, Y. and L. Li (2018). Matrix graph hypothesis testing and application in brain connectivity alternation detection. Statistica Sinica, to appear.

Zalesky, A., A. Fornito, and E. T. Bullmore (2010). Network-based statistic: identifying differences in brain networks. Neuroimage 53 (4), 1197–1207.

Zhang, T., Q. Yin, B. Caffo, Y. Sun, D. Boatman-Reich, et al. (2017). Bayesian inference of high-dimensional, cluster-structured ordinary differential equation models with applications to brain connectivity studies. The Annals of Applied Statistics 11 (2), 868–897.

